# Insight into the structural hierarchy of the protease cascade that regulates the mosquito melanization response

**DOI:** 10.1101/2023.07.13.548954

**Authors:** Sally A. Saab, Xiufeng Zhang, Suheir Zeineddine, Bianca Morejon, Kristin Michel, Mike A. Osta

## Abstract

Serine protease cascades regulate important insect immune responses, including melanization and Toll pathway activation. In the context of melanization, central components of these cascades are clip domain serine proteases (CLIPs) including the catalytic, <u>c</u>lip domain <u>s</u>erine <u>p</u>roteases (cSPs) and their non-catalytic <u>h</u>omologs (cSPHs). Here, we define partially the structural hierarchy of *An. gambiae* cSPs of the CLIPB family, central players in melanization, and characterize their relative contributions to bacterial melanization and to mosquito susceptibility to bacterial infections. Using *in vivo* genetic analysis we show that the protease cascade branches downstream of the cSPs CLIPB4 and CLIPB17 into two branches one converging on CLIPB10 and the second on CLIPB8. We also show that the contribution of key cSPHs to melanization *in vivo* in response to diverse microbial challenges is more significant than any of the individual cSPs, possibly due to partial functional redundancy among the latter. Interestingly, we show that the key cSPH CLIPA8 which is essential for the efficient activation cleavage of CLIPBs *in vivo* is efficiently cleaved itself by several CLIPBs *in vitro*, suggesting that cSPs and cSPHs regulate signal amplification and propagation in melanization cascades by providing positive reinforcement upstream and downstream of each other.

## Introduction

Clip domain serine proteases (CLIPs) function in cascades to regulate several insect immune responses, including particularly the activation of Toll signaling pathway [1, 2] and melanization [3, 4]. The CLIP gene family includes the catalytic (cSPs) and non-catalytic (cSPHs) CLIPs. Mosquitoes are attractive models to study these cascades due to the significant expansion of the CLIP gene family in their genomes [5, 6] and the propensity of cSPHs that play key roles in regulating these cascades [7–11]. The functional significance of the CLIP gene family expansion in mosquitoes is still unclear, but posits a more complex organization and regulation of these cascades. Both, cSPs and cSPHs are produced as zymogens that require proteolytic cleavage at a specific site in the N-terminus of the protease domain or in the clip domain, respectively, to become functional [2, 12–17]. Certain cSPHs, however, have predicted cleavage activation sites in the N-terminus of the protease-like domain, similar to cSPs [18].

CLIPs function in cascades whose roles in immunity have been mainly addressed in the context of the Toll pathway and melanization. These cascades have been well characterized in *Drosophila*, whereby microbial sensing activates a cascade of cSPs upstream of the Toll pathway [1, 19, 20]. The cascade branches downstream of the cSPs Hayan and Persephone to regulate also the fly’s melanization response [1]. In the tobacco hornworm *Manduca sexta*, the protease cascade branches downstream of HP21 into two axes; one axis culminates in the activation of <u>p</u>rophenoloxidase-<u>a</u>ctivating <u>p</u>rotease 2 (PAP2), PAP3 and two cSPHs (SPH1 and SPH2) leading to PPO activation, and the second culminates in the activation of HP6 which can further amplify the melanization response by cleaving PAP1, or activate Toll pathway by cleaving HP8, homolog of *Drosophila* Spe [14, 16, 17, 21, 22].

The genome of the malaria vector *An. gambiae* contains 110 CLIP genes, which are divided into 5 subfamilies; A, B, C, D and E [6, 18]. CLIPs B, C and D are predicted as catalytic cSPs, CLIPAs as non-catalytic cSPHs, whereas CLIPEs contain cSPHs or hybrid genes encoding proteins with both catalytic and non-catalytic protease domains. Genetic studies in *An. gambiae* identified several cSPHs as key regulators of melanization. SPCLIP1 (CLIPA30; [18]), CLIPA8 and CLIPA28 form a core cSPH module that is central to microbial melanization [7, 9, 10, 23]. These cSPHs exhibit a clear hierarchy of activation cleavage in the hemolymph with SPCLIP1 being most upstream, followed by CLIPA8 then CLIPA28 [7, 24]. So far, only the thioester-containing protein 1 (TEP1), a hallmark of mosquito humoral immunity [25], has been shown to act upstream of this cSPH module, controlling the activation cleavage of its constituent cSPHs [7, 24], whereas the identity of cSPs that cleave these cSPHs remain unknown. On the other hand, CLIPA2 and CLIPA14, serve as immune checkpoints that negatively regulate the mosquito melanization response to infection [8, 11]. Interestingly, the core cSPH module controls the cleavage of CLIPA14, possibly as a negative feedback loop to regulate the intensity of melanization [8].

Three experimental models for melanization, *P. berghei* ookinete melanization in *C-type lectin 4* (*CTL4*) knockdown (kd) mosquitoes, melanotic tumor formation in *serpin 2* (SRPN2) kd mosquitoes, and melanin quantification in mosquito excreta, have been used in genetic studies to identify several melanization-associated cSPs of the CLIPB and C families, including CLIPB4, B8, B9, B10, B14, B17 and CLIPC9 [23, 24, 26–28]. Surprisingly, none of these cSPs seems to regulate proteolysis within the core cSPH module *in vivo* [7, 24], raising questions about the molecular interactions that control cSPH activation cleavage in mosquito CLIP cascades, especially that lepidopteran cSPs, specifically PAPs, are known to cleave key cSPHs during the melanization response [15, 29, 30]. Unlike cSPHs, the organization of *An. gambiae* cSPs in the protease cascade regulating melanization remains largely unknown, with the exception of CLIPB4, CLIPB9 and CLIPB10 which function as PAPs, based on their ability to cleave *M. sexta* PPO *in vitro*, and CLIPB8 which seems to function upstream of CLIPB9 [26, 28, 31, 32]. Also, the individual contribution of these cSPs to the process of melanin formation in response to bacterial infections remains unexplored. Here, we define partially the hierarchical organization of *An. gambiae* cSPs involved in melanization and characterize their relative contributions to this immune reaction in response to different bacterial challenges. Using *in vivo* functional genetic analysis and *in vitro* assays, we also provide novel insight into the functional organization of cSPs and cSPHs in the protease cascade regulating mosquito melanization that advance our understanding of signal propagation and amplification in these cascades.

## 2. Materials and Methods

### 2.1. Experimental animal treatments and ethics of statement

This study was carried according to the recommendations in the Guide for the Care and Use of Laboratory Animals of the National Institutes of Health (Bethesda, USA). Animal protocol was approved by the Institutional Animal Care and Use committee IACUC of the American University of Beirut (permit number 18-08-504). The IACUC functions in compliance with the Public Health Service Policy on the Humane Care and Use of Laboratory Animals (USA), and adopts the Guide for the Care and Use of Laboratory Animals of the National Institutes of Health. Experiments were done using *An. gambiae* mosquito G3 strain. Mosquitoes were reared in the insectary at 27 °C (± 0.5) and 80% (± 5%) humidity with a 12 h day-night cycle as previously described [7].

### 2.2. Double Stranded RNA Synthesis and Gene Silencing

DNA amplification of the genes of interest was performed using the T7 flanked primers listed in Table S1. <u>D</u>ouble <u>s</u>tranded RNA (dsRNA) was synthesized from purified T7-tagged PCR amplicons using the T7 RiboMax Express Large-Scale RNA production system (Promega) according to the manufacturer’s instructions and purified as previously described [33]. *In vivo* gene silencing by RNAi was performed as previously described [33]. Briefly, 1 to 3-days-old female adult mosquitoes were microinjected intrathoracically with 69 nl of a 3.5 μg/μl gene-specific dsRNA solution in water under CO2, using a Drummond Nanoject II nanoliter injector. Mosquitoes were allowed 2-4 days to recover before further manipulation. To determine whether cSPs exhibit redundancy in cSPH cleavage, we scored the effect of simultaneously silencing CLIPB4, B8, B9, B10, B17 and CLIPC9 on the cleavage of SPCLIP1 and CLIPA28 in the hemolymph at 1 hour after *S. aureus* (OD600=1) injection. The simultaneous silencing of all six genes was done as follows: Freshly emerged mosquitoes were injected each with 138 nl of a dsRNA mixture containing 5 μg/μl of each of ds*CLIPB4*, ds*CLIPB9*, ds*CLIPB10* and ds*CLIPB17*, then mosquitoes were allowed to rest for 2 days before receiving 138 nl injection of another dsRNA mixture containing 3.5 μg/μl of each of ds*CLIPB8* and ds*CLIPC9*. We adopted this 2-step strategy to silence all 6 genes since we could not deliver all 6 dsRNAs in one injection which would render the solution too viscous to be injected with our Nanoinjector.

Efficiency of gene silencing was measured by western blot analysis where antibodies are available, as follows. Hemolymph was extracted from approximately 40 naïve mosquitoes per genotype including, the ds*LacZ* control, at 3 to 4 days after dsRNA injection and proteins were quantified using Bradford Reagent (Fermentas). Equal amounts of proteins were resolved by reducing SDS-PAGE, transferred to Immuno-blot PVDF membrane (Bio-Rad) using wet transfer (Bio-Rad) and subjected to western blot analysis with the appropriate antibody dilution (western blot section below). The gene silencing efficiency of *Rel1*, *CLIPB14* and *CLIPB17* was measured by qRT-PCR. Total nucleic acids were extracted from 15 mosquitoes per genotype, including ds*LacZ*, using TRIzol reagent (Invitrogen). First strand cDNA synthesis and qRT-PCR were performed as previously described [34]. Primers used in qRT-PCR are listed in Table S2. Relative gene expression values were calculated using the comparative C_T_ method.

### 2.3. Hemolymph cleavage profile of cSPs by western blot analysis

Wild type or dsRNA treated adult *An. gambiae* female mosquitoes were infected with the GFP-expressing, ampicillin-resistant *Escherichia coli* OP-50 strain (OD_600_= 0.8 or 3) [35], the tetracycline resistant *Staphylococcus aureus* (OD_600_= 0.8 or 1.5) [10] or *Beauveria bassiana* spores suspended in water (2000 spores/mosquito), prepared as described previously [36]. Hemolymph was extracted by proboscis clipping from groups of 40 mosquitoes per genotype in sterile 1x phosphate-buffered saline (PBS) containing a protease inhibitor cocktail (Roche), and protein concentration determined using Bradford protein quantification assay (Fermentas). Equal amounts of protein from the different hemolymph samples were resolved by SDS-PAGE and wet transferred to immunoblot PVDF membranes using BioRad apparatus. Primary antibodies used in western blotting include, rabbit αCLIPB4 (1:3000; produced by BOSTER), rabbit αCLIPB8 (1:1000;[31]), rabbit αCLIPB9 (1:1000; produced by BOSTER), rabbit αCLIPB10 (1:1000; [28]), rabbit αCLIPC9 (1:2000; [24]), rabbit αCLIPA2 (1:1000; [11]), rabbit αCLIPA14 (1:3000, produced by BOSTER), mouse αCLIPA8 (1:100; [10]), rabbit αCLIPA28 (1:1000; [7]), rabbit αSPCLIP1 (1:2000; produced by BOSTER), rabbit αTEP1 (1:1000; [7]), rabbit αSRPN2 (1:1000; [37]), mouse αApolipophorin II (1:100; [38]) and αCTL4 (1:1000; produced by Boster). Anti-rabbit or anti-mouse IgG horseradish peroxidase-conjugated secondary antibodies were used at dilutions of 1:15000 or 1:6000, respectively. Western blots were revealed with BioRad Clarity Max western ECL substrate and imaged using ChemiDoc MP (BioRad).

### 2.4. Melanization-Associated Spot Assay (MelASA)

DsRNA-treated mosquitoes were injected with *Escherichia coli* OP-50 strain (OD600 =3; [35]), tetracycline resistant *Staphylococcus aureus* strain (OD600 =3; [10]), and lyophilized *M. luteus* (ATCC, No. 4698, MilliporeSigma) resuspended in 1x PBS to an OD600=5 or with *B. bassiana* spores (100000 spores/mosquito) resuspended in water. Groups of 35 (for *E. coli* and *S. aureus* and *B. bassiana* infections) or 50 (for *M. luteus* infections) infected mosquitoes per genotype were housed in carton cups containing clean, white Whatman filter papers (5.5 cm diameter) at the bottom. Twelve (for *E. coli* and *S. aureus*), 14 (for *B. bassiana*) and 18 hours (for *M. luteus*) post-infections, the filter papers containing melanin spots in mosquito fecal excretions were imaged under white epi-illumination without filters in a ChemiDoc MP (BioRad). Image processing and total spot area calculations were performed as described previously [24]. Experiments were repeated at least six times. Differences in Total melanotic spot area were analyzed using unpaired t-tests to compare two treatment groups, and One-Way ANOVA with Dunnett’s post-test to compare multiple treatment groups in GraphPad Prism (version 9.5.1). Means were considered to differ significantly if *P* < 0.05.

### 2.5. Bacterial Proliferation and Mosquito Survival Assays

Mosquito resistance to bacterial infections was determined by scoring the Colony Forming Units (CFU) counts in whole homogenates of dsRNA-treated mosquitoes injected intrathoracically with the *E. coli* OP-50 strain or the tetracycline resistant *S. aureus* strain suspended in 1x PBS at an OD_600_=0.8, as previously described [7, 39]. Results from at least five different experiments were analyzed using the Kruskal-Wallis test followed by the Dunn’s multiple comparison test that compares mean ranks of each group to the ds*LacZ* control. Means of different groups were considered significantly different if the *P* <0.05.

To score mosquito survival following bacterial challenge, dsRNA-treated mosquitoes were injected with *E. coli* or *S. aureus* strains at OD_600_= 0.8 and OD_600_= 2, respectively. DsLacZ mosquitoes were used a negative control, while ds*CTL4* and ds*Rel1* mosquitoes were used as positive controls for *E. coli* and *S. aureus* infections, respectively. Mosquito survival was scored daily over two weeks. The Kaplan-Meier survival test was used to calculate percent survival. Experiments were repeated at least five times and the statistical significance of the observed differences was calculated using the Log-rank test. One representative experiment is shown.

### 2.6. In vitro assays to study CLIPA8 cleavage by candidate CLIPBs

To construct the CLIPA8 expression vector, a sequence encoding Asp-Leu-V5 tag (GKPIPNPLLGLDST)- Arg- Thr- Gly-His tag (HHHHHH)-stop codon was inserted into the BglII and PacI sites in pOET3. The full-length CLIPA8 coding sequence, amplified using primers CLIPA8-F-NotI and CLIPA8-R-BglII, was cloned subsequently into NotI and BglII sites of the same plasmid to generate pOET3-CLIPA8-V5-His (for primer sequences see Table S3). The predicted CLIPA8 cleavage site ^63^IMLR^66^ was replaced by IEGR [26] using the CLIPA8mutag primer and the QuikChange Multi Site-Directed Mutagenesis Kit (Agilent Technologies) to create plasmid pOET3-CLIPA8_Xa_-V5-His (Table S3). P0 viruses were generated with FlashBAC Gold - Baculovirus Expression System (Genway Biotech) in Sf9 cells. For protein expression, a 800 ml Sf-900II suspension culture (2×10^6^ cells/ml) was inoculated at an MOI of 1 with P1 or P2 viruses, and media were harvested 2 days later. Protein purifications were performed as described previously, using sequentially dialysis, Ni-NTA agarose columns (Qiagen), and Q Sepharose Fast Flow columns (Cytiva) [28, 31]. Recombinant protein was concentrated and buffer exchanged to 20 mM Tris•HCl, 50 mM NaCl, pH 8.0 using Amicon Ultra centrifugal filter units (Sigma-Aldrich). Protein purity was confirmed by CBB staining and Western blot analyses using anti- THE^TM^ His Tag Antibody, THE^TM^ V5 Tag antibody (both at 1:5000 dilution, GenScript), and CLIPA8 primary antibody (1:5000 dilution, Boster Biological Technology). To study the cleavage of CLIPA8 by candidate CLIPBs, Factor Xa activation of proCLIPB4_Xa_-His, proCLIPB9_Xa_-His, and proCLIPB10_Xa_-His was performed as described previously [26, 28, 31]. To cleave proCLIPA8_Xa_-V5-His by Factor Xa, 4 µg of proCLIPA8_Xa_ was incubated at RT for 2 hours with 1 µg of Factor Xa in a total volume of 40 µl activation buffer (20 mM Tris•HCl, 100 mM NaCl, 2mM CaCl_2_, pH 8.0). To assess the cleavage pattern of wild-type proCLIPA8-V5-His by B4, B9 and B10, 1 µl of proCLIPA8 (250 ng/µl) was mixed with 375 ng Factor Xa-activated B9_Xa_, 200 ng Factor Xa-activated B4_Xa,_ and 10 ng Factor Xa-activated B10_Xa_, respectively and incubated in reaction buffer (20 mM Tris•HCl, 80 mM NaCl, 2.25 mM CaCl_2_, pH 7.5) in a 10 µl total reaction volume, either on ice for 30 min (B10_Xa_) or at RT for 1 h (B4_Xa_ and B9_Xa_). CLIPA8 cleavage was assessed by subjecting reactions to Western blot analyses using anti-His and anti-V5 antibodies.

## 3. Results

### 3.1. Systemic infections trigger the cleavage of CLIPB10 and consumption of CLIPB8

In *An. gambiae*, out of the 63 cSPs identified in the genome [6, 18], several CLIPB members including, CLIPB4, CLIPB8, CLIPB9, CLIPB10, CLIPB14, CLIPB17 were shown to play a significant role in *P. berghei* melanization in *CTL4* kd refractory mosquitoes and in melanotic tumor formation in naïve *SRPN2* kd mosquitoes [23, 24, 26–28]. To build hierarchy among CLIPBs, we characterized first their hemolymph cleavage profiles in response to infection as readouts for subsequent genetic epistasis analyses, as previously described for cSPHs [7, 24]. No hemolymph cleavage products were detected for CLIPB4 and CLIPB9 after septic infections with *E. coli*, *S. aureus* or *B. bassiana* (Fig. S1). With respect to CLIPB8, its ∼48kDa full-length form detected in the hemolymph of naïve mosquitoes was consumed completely at 24 and 10 hrs following *B. bassiana* (Fig.1 A) and *S. aureus* infections (Fig.1 B), respectively, but was not altered by *E. coli* infections (Fig.1 C). Full-length CLIPB10 (CLIPB10-F) of ∼ 40 kDa was cleaved at later time points after *S. aureus* infections resulting in a weak, lower molecular weight band (CLIPB10-C) of ∼35 kDa (Fig. 1 E) corresponding to the C-terminal protease domain (antibody was raised against the protease domain [28]). CLIPB10 cleavage was not detected after *B. bassiana* (Fig. 1 D) nor *E. coli* (Fig. 1 F) infections, suggesting that Gram-positive bacteria constitute a stronger stimulus for CLIPB10 cleavage. Unfortunately, we were not able to characterize CLIPB17 and CLIPB14 proteins since their custom-made antibodies failed to detect specific bands in the hemolymph.

**Fig. 1.**
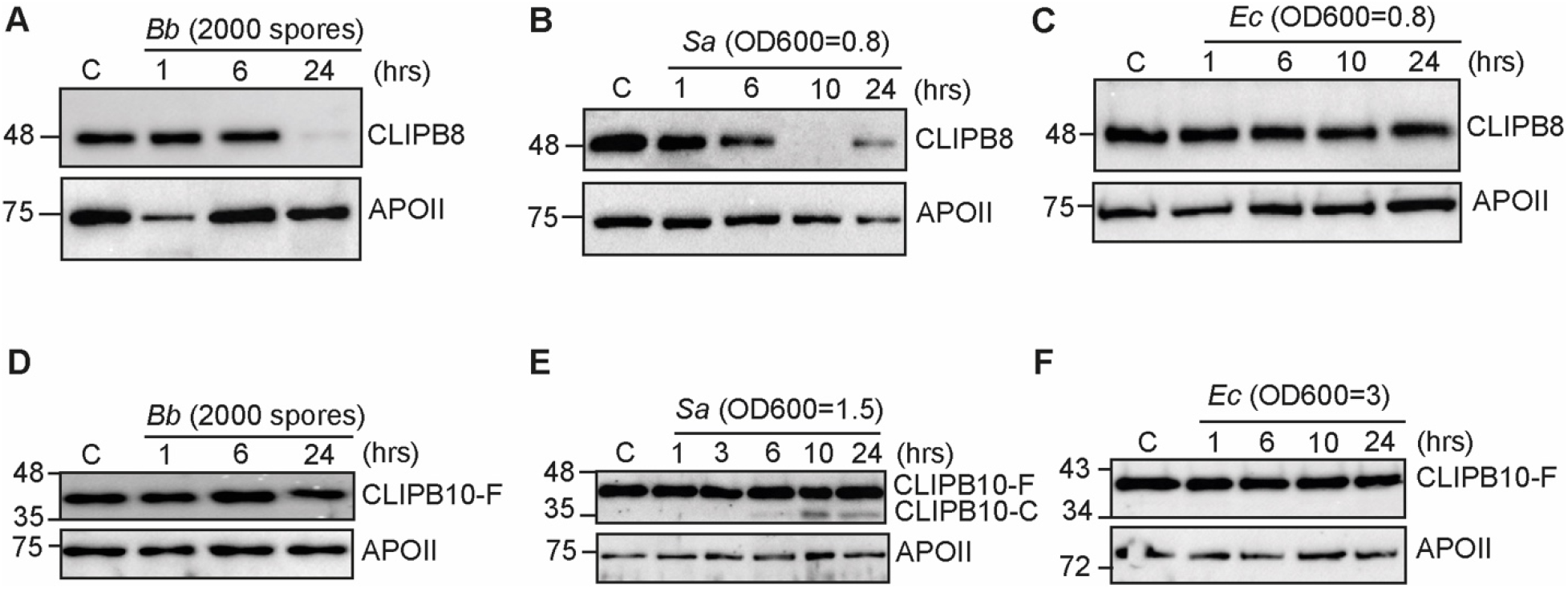
CLIPB8 and CLIPB10 hemolymph proteins respond to microbial infection. CLIPB8 is consumed in the hemolymph in response to (A) *B. bassiana* and (B) *S. aureus*, but not (C) *E. coli* infection. CLIPB10 is cleaved at 24 hours after (E) *S. aureus* but not (D) *B. bassiana* nor (F) *E. coli* infections. Western blots of hemolymph extracts obtained at the indicated time points from (A,D) *B. bassiana* (2000 spores/mosquito), (B,E) *S. aureus* and (C,F) *E. coli* injected mosquitoes and probed with αCLIPB8. Western blots were probed with (A-C) αCLIPB8 and (D-F) αCLIPB10, respectively. All hemolymph samples were extracted from 40 mosquitoes, quantified using Bradford protein quantification assay, and equal amounts of protein were loaded into each well. Membranes were reprobed with αAPOII (without stripping) to control for loading. The figures shown are representative of 3 independent biological experiments. *C*, control naïve mosquitoes.

### 3.2. CLIPB4 and CLIPB17 act upstream of CLIPB8 and CLIPB10 in the protease cascade regulating melanization

We utilized CLIPB8 depletion and CLIPB10 cleavage patterns in the hemolymph of *S. aureus* infected mosquitoes as readouts to assign hierarchy among the CLIPBs involved in melanization. Silencing *CLIPB4* and *CLIPB17* rescued CLIPB8 depletion (Fig. 2 A) and abolished CLIPB10 cleavage (Fig. 2 B) in the hemolymph at 10 and 24 hrs after *S. aureus* infection, respectively (Fig. 2 A), indicating that CLIPB4 and CLIPB17 act upstream of CLIPB8 and CLIPB10. The fact that *CLIPB8* kd did not abolish CLIPB10 cleavage and that *CLIPB10* kd did not rescue CLIPB8 depletion suggests that the cascade bifurcates downstream of CLIPB4 and CLIPB17 into two branches; one converging on CLIPB8 and the other on CLIPB10 (Fig. 2 C). Interestingly, *SPCLIP1* silencing rescued CLIPB8 depletion (Fig. 2 A) and abolished CLIPB10 cleavage (Fig. 2 B), further confirming its upstream position in the melanization response as part of the core [SPCLIP1-CLIPA8-CLIPA28] cSPH module [7, 24]. CLIPB8 silencing almost depleted CLIPB8 from the hemolymph (Fig. 2 A), whereas CLIPB10 silencing significantly reduced CLIPB10 hemolymph levels, suggesting that CLIPB10 is either abundantly produced or that certain cell populations that express CLIPB10 show some resistance to RNAi. The silencing efficiencies of the remaining genes used in this assay are shown in Fig. S2.

**Fig. 2.**
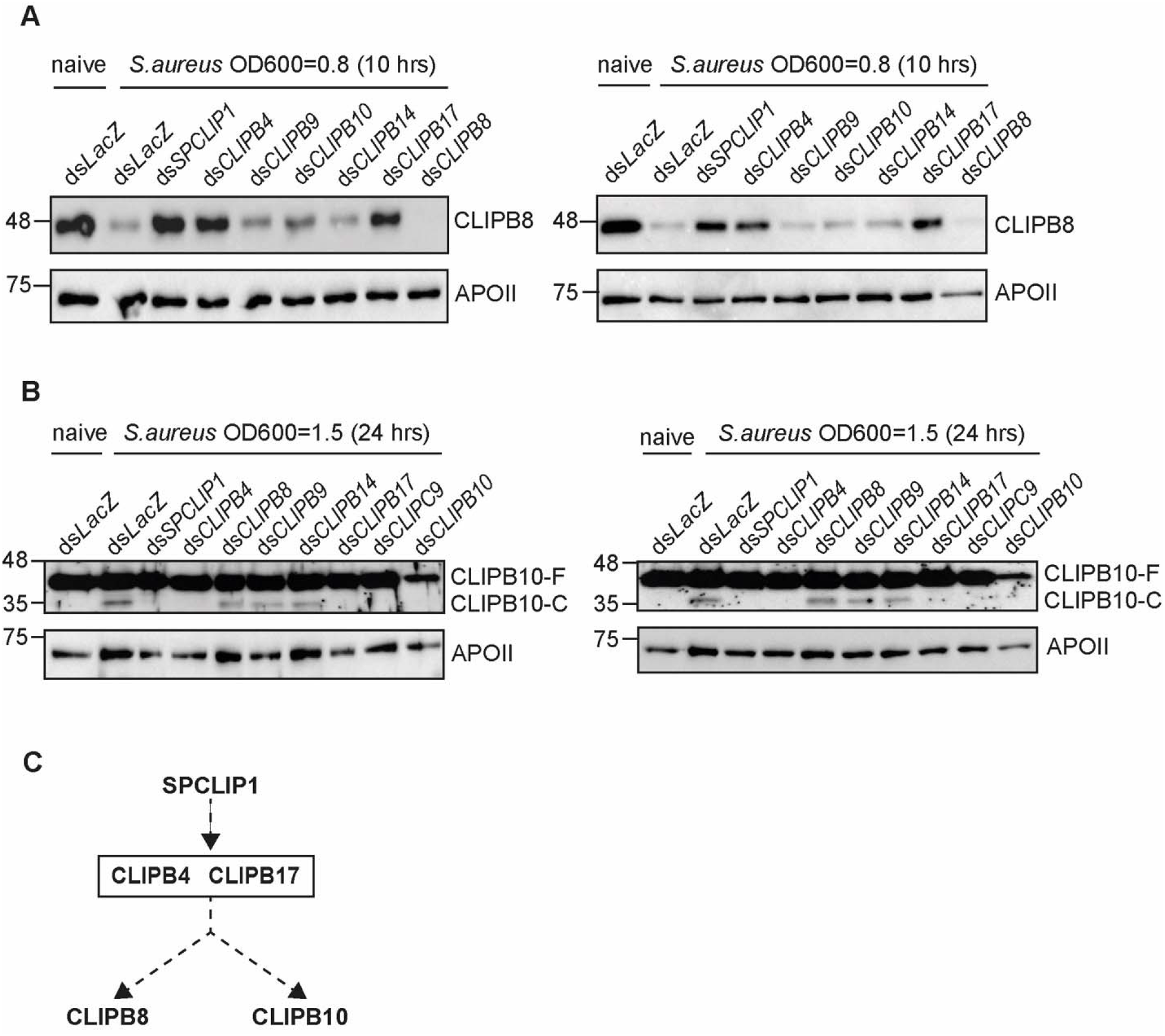
CLIPB4 and CLIPB17 act upstream of CLIPB8 and CLIPB10 in the serine protease cascade regulating melanization. Western blots showing (A) CLIPB8 protein levels and (B) CLIPB10 cleavage profile in the hemolymph of mosquitoes treated with the indicated gene-specific double-stranded (ds) RNA at 10 and 24 hrs after *S. aureus* infection, respectively. Two representative trials are shown per experiment. All hemolymph samples were extracted from 40 mosquitoes, quantified using Bradford protein quantification assay, and equal protein amounts were loaded into each well. Membranes were reprobed with αAPOII (without stripping) as loading control. (C) Schematic diagram showing the hierarchical organization of SPCLIP1 with respect to the catalytic CLIPBs in the melanization response to systemic bacterial infections. CLIPB4 and CLIPB17 are shown in a rectangular box because their respective hierarchies have not been determined. Dashed lines indicate that the enzymatic steps are not yet fully characterized.

### 3.3. TEP1 and the core cSPH module are upstream regulators of CLIPBs in the melanization response

The fact that SPCLIP1 controlled the activation of CLIPB8 and CLIPB10, prompted us to test whether the other two members of the cSPH module exhibit a similar phenotype. Indeed, silencing either CLIPA8 or CLIPA28 abolished CLIPB10 cleavage (Fig. 3 A) and rescued CLIPB8 depletion (Fig. 3 B) in the hemolymph of *S. aureus* infected mosquitoes. These results, together with the reported role of the core cSPH module as upstream regulator of CLIPC9 [24], strongly suggest that this module acts, upstream of all catalytic CLIPs with reported roles in *An. gambiae* melanization (Fig. 3 C).

**Fig. 3.**
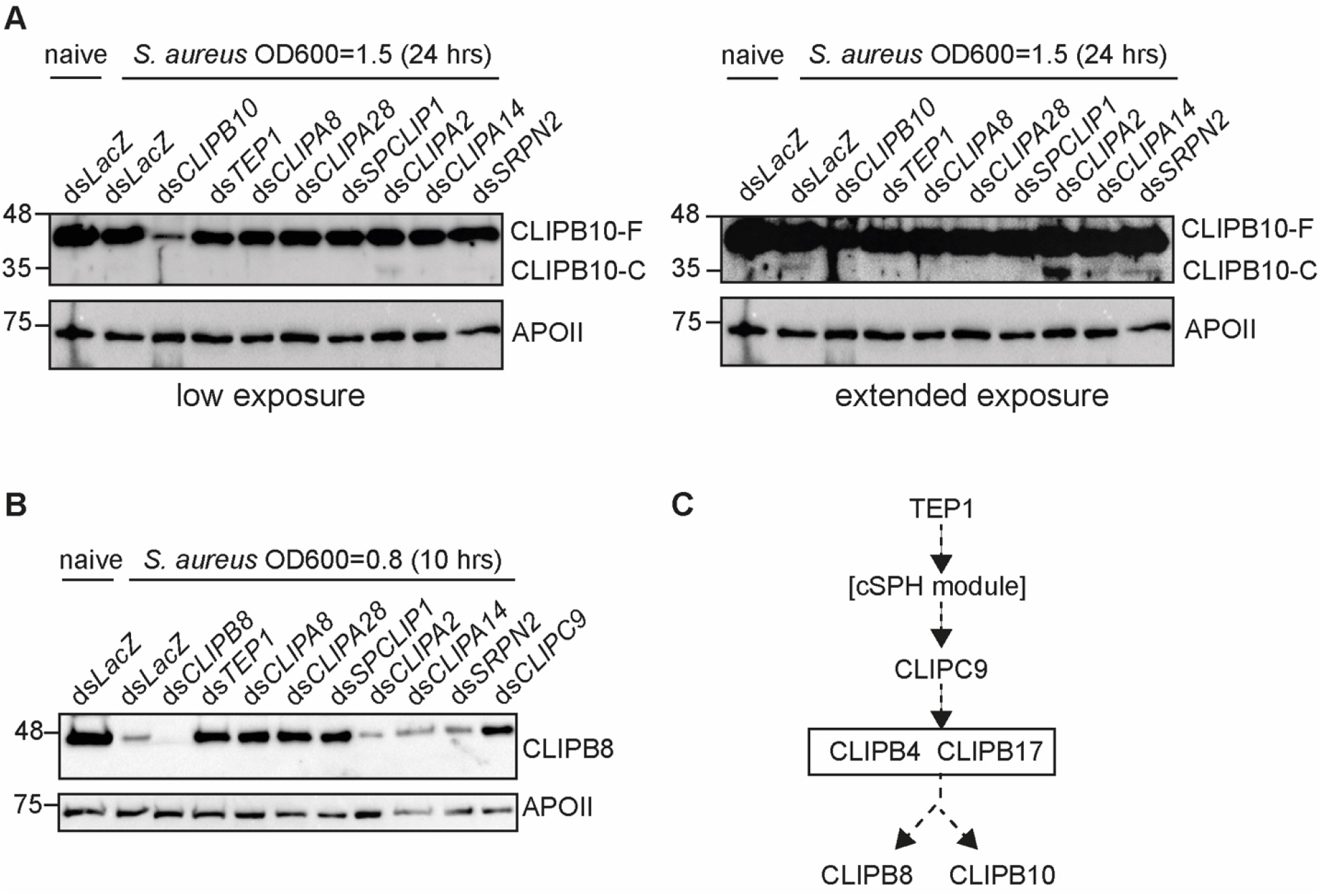
TEP1 and the core cSPH module control the activation of CLIPB8 and CLIPB10. Western blots showing (A) CLIPB10 cleavage profile and (B) CLIPB8 protein levels in the hemolymph of mosquitoes treated with the indicated gene-specific double-stranded (ds) RNAs at 10 and 24 hrs after *S. aureus* infection, respectively. In (A), low (left) and extended (right) exposures of the same membrane are shown in order to reveal the CLIPB10-C band that is often very weak. All hemolymph samples were extracted from 40 mosquitoes, quantified using Bradford protein quantification assay, and equal protein amounts were loaded into each well. Membranes were reprobed with αAPOII (without stripping) as loading control. The figures shown are representative of 3 independent biological experiments. (C) Schematic diagram showing the hierarchical organization of TEP1, the core cSPH module and the catalytic CLIPBs and CLIPC9 in the melanization response to systemic bacterial infections. Dashed lines indicate that the enzymatic steps are not yet fully characterized.

Only TEP1 has been shown to control proteolytic activation of the core cSPH module [7, 9]. Hence, we asked whether TEP1 also regulates CLIPB8 and CLIPB10 activation cleavage during systemic infections. Indeed, TEP1 silencing abolished CLIPB10 cleavage (Fig. 3 A) and rescued CLIPB8 depletion (Fig. 3 B) in the hemolymph of *S. aureus* infected mosquitoes, placing TEP1 as the most upstream factor in the melanization response (Fig. 3 C).

CLIPC9 functions downstream of the core cSPH module in the melanization response [24]. Here, we show that *CLIPC9* kd rescued CLIPB8 depletion (Fig. 3 B) and abolished CLIPB10 cleavage (Fig. 2 B) after *S. aureus* infections, suggesting that it acts upstream of these two CLIPBs. Although CLIPCs are known to function upstream of CLIPBs in the protease cascades (reviewed in [3]), it would be interesting to determine whether CLIPC9 activation cleavage is subject to positive feedback or reinforcement by downstream members of the CLIPB family.

CLIPA2, CLIPA14 and SRPN2 are potent negative regulators of the melanization response [7, 8, 11, 23, 24, 31], yet silencing neither of these negative regulators influenced CLIPB10 cleavage (Fig. 3 A) and CLIPB8 consumption (Fig. 3 B) patterns in the hemolymph. Although *CLIPA2* kd appears to enhance CLIPB10 cleavage, this phenotype was not consistently observed (data not shown). There are multiple equally parsimonious explanations for the observations, including that the intervention points for CLIPA2 and A14 are downstream of CLIPB8 and B10, and that SRPN2 in not an inhibitor of the protease that cleaves CLIPB10.

### 3.4. CLIPA8 is cleaved by several CLIPBs in vitro

It is surprising that only TEP1 silencing, so far, controls the proteolytic processing of members of the core cSPH module in response to systemic microbial infections [7, 9], whereas, silencing none of the key cSPs involved in mosquito melanization was able to phenocopy TEP1 [7, 24]. We hypothesize that several cSPs may cleave the same cSPH to trigger the rapid activation of these essential factors, which could explain why single cSP gene knockdowns failed to rescue cSPH cleavage during systemic infections [7, 24]. To address this question, we characterized the cleavage pattern of CLIPA8 *in vitro*. We chose CLIPA8 for the following reasons: (1) CLIPA8 is a member of the cSPH module that controls melanization in *An. gambiae*. (2) *SRPN2* kd enhances the cleavage of full-length CLIPA8 *in vivo* [7], and (3) SRPN2 directly inhibits the protease activities of CLIPB4, B9 and B10 [26, 28, 32], rendering these three proteases as potential candidates for CLIPA8 cleavage. To that purpose, we expressed two versions of full-length CLIPA8; a wildtype version containing two C-terminal tags, V5 and His (CLIPA8-V5-His) and a mutant version in which the predicted cleavage site IMLR (cleavage predicted to occur after the R residue [18]) was replaced with IEGR (CLIPA8_Xa_-V5-His) to allow cleavage by commercial Factor Xa. Both versions were obtained with high purity (minimally 95%) at ∼ 54 kDa, and reacted with antibodies against CLIPA8, V5 and His, as expected (Fig. S3). The theoretical weight of recombinant CLIPA8 is ∼40 kDa based on the constructed plasmid pOET3-CLIPA8-V5-His, suggesting that the observed size shift is likely caused by protein modifications in the eukaryotic cell expression system.

Zymogens proCLIPB4_Xa_-His, proCLIPB9_Xa_-His and proCLIPB10_Xa_-His containing a C-terminal His were expressed and purified as previously described [26, 28, 31]. Factor Xa was used to activate these three zymogens, and cleaved products corresponding to active CLIPB4_Xa_ (Fig. 4 A), CLIPB9_Xa_ (Fig. 4 B) and CLIB10_Xa_ (Fig. 4 C) were detected by western blot analyses using anti-His antibody. *In vitro* assays revealed that CLIPA8-V5-His was efficiently cleaved by Factor Xa-activated CLIPB4_Xa_ (Fig. 4 A), CLIPB9_Xa_ (Fig. 4 B) and CLIB10_Xa_ (Fig. 4 C). Cleaved CLIPA8 was detected at the same size as that of cleaved CLIPA8_Xa_, suggesting that CLIPB4, B9 and B10 likely cleave CLIPA8 after the R residue in IMLR. Minimal cleavage of CLIPA8 was observed when incubated alone with proCLIPB4_Xa_-His (Fig. 4 A) or proCLIPB10_Xa_-His (Fig. 4 C), suggesting these CLIPB zymogens might have been minimally activated during purification. Altogether, our results suggest that cSPs may exhibit redundancy with respect to the cleavage of cSPHs in the core module. To determine whether this redundancy in cSPH cleavage is also observed *in vivo*, we monitored the cleavage profiles of CLIPA28 (which is downstream of CLIPA8 in the cSPH module) and SPCLIP1 (which is upstream of CLIPA8 in the cSPH module) in *S. aureus*-injected mosquitoes in which CLIPB4, B8, B9, B10, B17 and CLIPC9 were silenced simultaneously. This multi-gene silencing approach triggered efficient gene silencing, as shown for CLIPB10 and CLIPB4 as selected indicators (Fig. S4), that is similar to what we observe with the individual gene knockdowns. The results showed that CLIPA28 and SPCLIP1 were still efficiently cleaved in these mosquitoes (Fig. S4). A plausible explanation for these results is that the residual enzymatic activities of the silenced cSPs, due to incomplete gene silencing by RNAi (as shown in Fig. S4 for CLIPB10 and CLIPB4), are still sufficient to efficiently cleave the cSPHs. Alternatively, other yet to be discovered cSPs may also be contributing the cSPH cleavage. Nevertheless, these results support the *in vitro* observations that cSPHs are likely to be cleaved by more than one cSP.

**Fig. 4.**
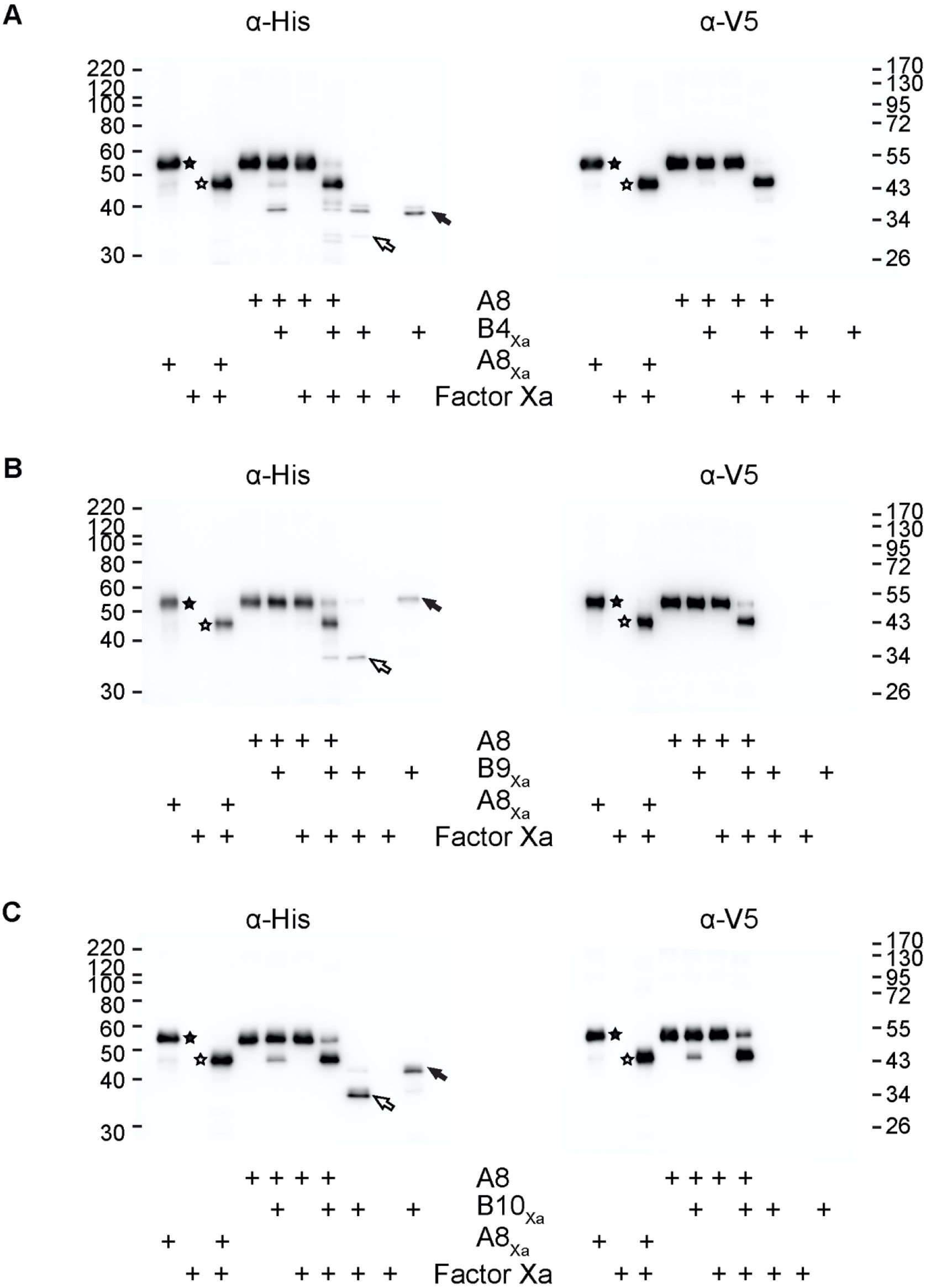
Recombinant proCLIPA8 is cleaved by B4, B9 and B10 *in vitro*. Recombinant proCLIPA8-V5-His is cleaved by (A) Factor Xa-activated B4_Xa_-His, (B) B9_Xa_-His, and (C) B10_Xa_-His at the predicted site, examined by Western blot analyses using anti-His and anti-V5 antibodies. Zymogens of CLIPB members were activated by Factor Xa, and incubated with wild-type proCLIPA8 for (A,B) 1 hour at RT or (C) 30 min on ice. The activation of corresponding CLIPB members by Factor Xa was included for each Western blot analysis. A positive control was included with Factor Xa-cleavage of proCLIPA8_Xa_ for 2 hours at RT. Both anti-His and anti-V5 antibodies detected the cleaved fragment of wild-type proCLIPA8 at a similar size as that of Factor Xa-cleaved proCLIPA8_Xa_. Solid and hollow arrows indicate the full-length and activated forms of CLIPB members, respectively; for CLIPA8_Xa_, solid and hollow stars indicate the full-length and cleaved forms, respectively.

### 3.5. CLIPB4 and CLIPB17 are central players in the melanization response to Gram-negative and Gram-positive bacteria

CLIPB4, CLIPB8, CLIPB9, CLIPB10, CLIPB14 and CLIPB17 are involved in melanotic tumor formation and/or *P. berghei* melanization [23, 26–28, 31], however, their implication in bacterial melanization have not been addressed yet. This is especially important, given the more significant evolutionary pressure that is likely to be imposed by bacteria on the mosquito immune system, especially in their larval habitats. In contrast, only a small percentage of mosquitoes in the field are infected with *Plasmodium* parasites and there is no evidence that mosquito immunity genes have been subject to selection by malaria parasites [40, 41]. To gauge the contribution of the candidate CLIPBs in bacterial melanization, we used the Melanization-associated Spot Assay (MelASA) [24]. Silencing CLIPB4 and CLIPB17 but not CLIPB8, CLIPB9, CLIPB10 or CLIPB14 significantly reduced the total spot area of melanin deposits at 12 hrs after mosquito injections with low (OD_600_=0.8) or high (OD_600_=3) doses of *E. coli* compared to ds*LacZ* controls (Fig. 5 A). However, the knockdown of neither of these CLIPBs reduced the total melanin spot area after *S. aureus* infections (Fig. 5 B). To determine whether the phenotype observed with *S. aureus* spans other Gram-positives, we challenged mosquitoes with *M. luteus*, a low-virulence Gram-positive bacteria widely used in insect immune challenges [20, 42, 43] and known to be a potent trigger of mosquito melanization [42, 44]. All tested CLIPBs, except CLIPB14, showed significant contribution to the melanization response triggered by *M. luteus* (Fig. 5 C). On the other hand, *SPCLIP1* silencing almost abolished melanin excreta in response to all bacterial infections (Fig. 5), further supporting the essential role the core cSPH module plays in the melanization response. However, neither SPCLIP1 knockdown nor that of any of the tested CLIPBs significantly reduced melanin excreta in mosquitoes after 14 hours of *B. bassiana* spore injections (Fig. S5). These results are not surprising, and are in line with our previous study showing that the melanization response to the early stages of *B. bassiana* development in the mosquito is hemocyte-mediated and independent of TEP1 and CLIPA8 [45].

**Fig. 5.**
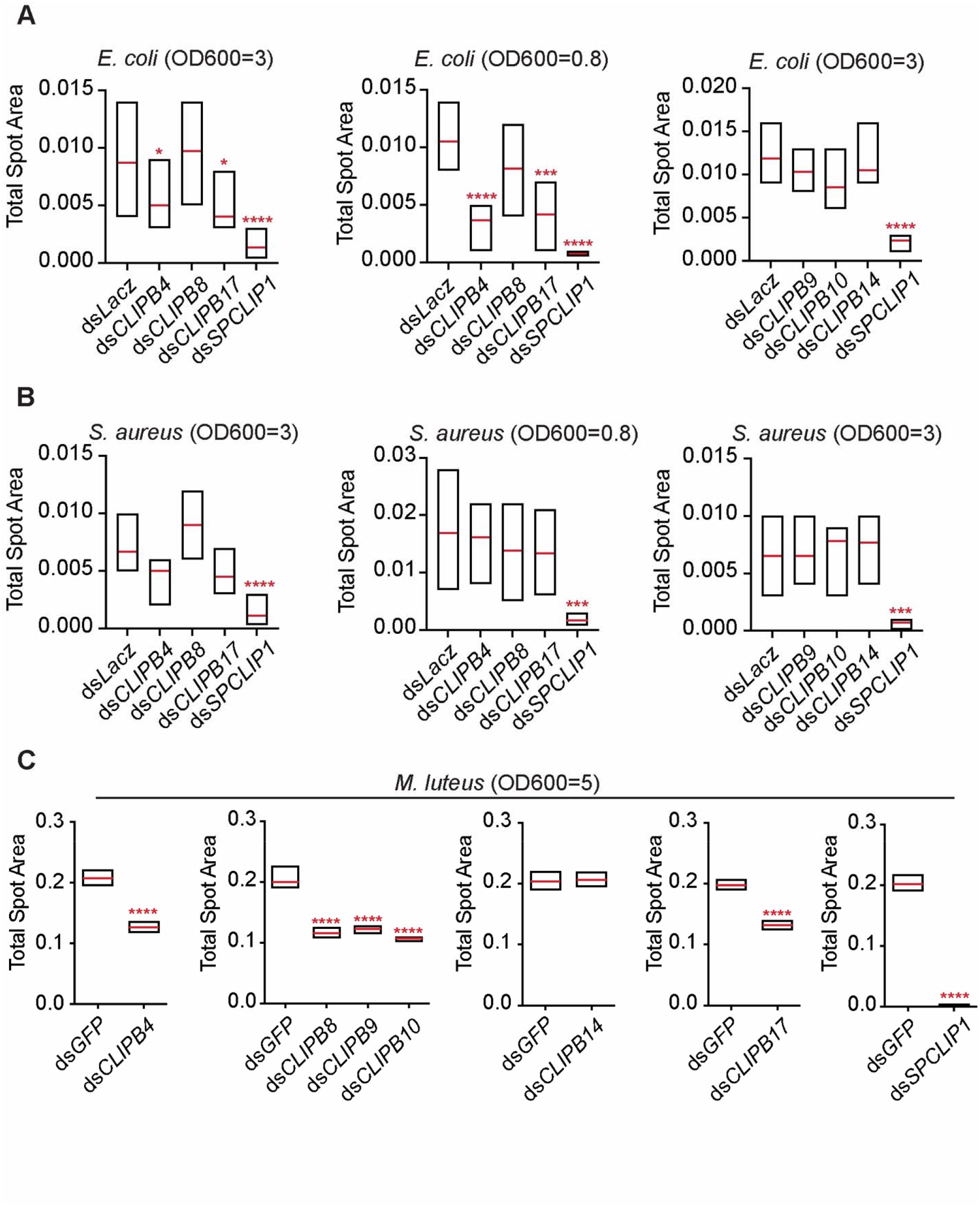
CLIPB4 and CLIPB17 are central players in the melanization response to bacterial infections. MelASA assay conducted on the indicated mosquito genotypes at 12 hrs post (A) *E. coli*, (B) *S. aureus* and 18 hrs post (C) *M. luteus* infections. The amount of melanin deposits present in infected mosquito excreta was measured from Whatman papers placed at the bottom of the paper cups housing 35 (for *E. coli* and *S. aureus*) or 50 (for *M. luteus*) mosquitoes per sample. Statistical analysis was done using unpaired t-tests to compare two treatment groups, and One-Way ANOVA with Dunnett’s post-test to compare multiple treatment groups. Means are shown in red lines. Data shown are from at least 6 independent trials. *, *P*<0.05; ***, *P*<0.001; ****, *P*<0.0001.

### 3.6. CLIPB14 regulates mosquito tolerance but not resistance to bacterial infections

Certain cSPs function at the crossroads of PPO and Toll pathway activation [1, 17, 21], and silencing key immune-related cSPs in *Drosophila* increased the fly susceptibility to microbial infections [1, 46, 47]. We therefore tested whether any of our candidate CLIPBs contribute to mosquito tolerance and/or resistance to systemic bacterial infections. Tolerance, an indicator of host fitness to infection, was measured by scoring the survival of ds*CLIPB* mosquitoes after intrathoracic injections with *S. aureus* or *E. coli*. Ds*CTL4* [39, 48] and ds*Rel1* [49] mosquitoes were used as positive controls for *E. coli* and *S. aureus* infections, respectively. Only ds*CLIPB14* mosquitoes showed compromised survival to infections with *E. coli* (Fig. 6 C) and *S. aureus* (Fig. 6 D), whereas silencing the other CLIPBs had no effect (Fig. 6 A-D). Of note, the compromised survival of ds*CLIPB14* mosquitoes was only observed at low (OD_600_=0.8) but not high (OD_600_=2) infection doses. We utilized two different infection doses in these survival assays, since a previous study in *Drosophila* showed that the fly’s survival to low and high doses of *S. aureus* infections relies on distinct immune responses [1]. Mosquito resistance, an indicator of immune-mediated microbial clearance, to *E. coli* and *S. aureus* systemic infections was measured by scoring the CFUs in whole mosquito homogenates. Interestingly, silencing neither of the CLIPBs influenced the ability of mosquitoes to clear either bacteria from the tissues (Fig. 6 E-F). The same phenotype was also observed for *Rel1* kd, which is in line with our previous study [49], suggesting that Toll pathway might not be essential to *S. aureus* clearance at that time point post-infection.

**Fig. 6.**
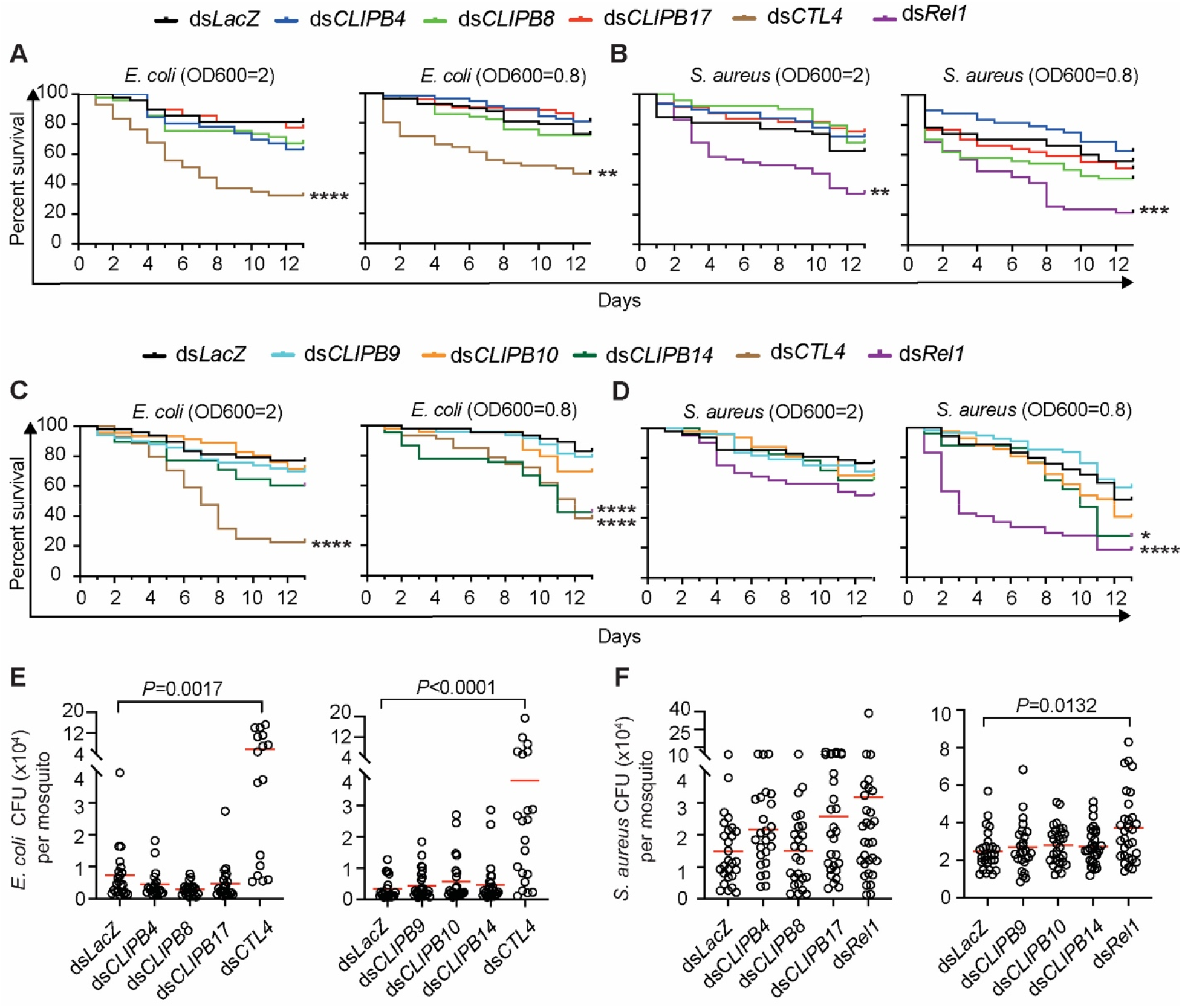
CLIPB14 is required for mosquito tolerance but not resistance to bacterial infections. (A-D) Survival analysis of the indicated mosquito genotypes challenged with (A and C) *E. coli*, and (B and D) *S. aureus*. A representative experiment of at least three independent trials is shown. The Kaplan-Meier survival test was used to calculate the percent survival. Statistical significance of the observed differences was calculated using the Log-rank test. *, *P*<0.05; **, *P*<0.01; ***, *P*<0.001; ****, *P*<0.0001. (E, F) Bacterial proliferation assays conducted on the indicated mosquito genotypes injected with (E) *E. coli* (OD_600_=0.8) and (F) *S. aureus* (OD_600_=0.8). Batches of 8 whole mosquitoes each were homogenized in LB medium at 24 hours post-infection and colony forming units (CFUs) were scored on LB plates supplemented with the appropriate antibiotic. Each point on the scatter plot represents the mean CFU per mosquito per batch. Statistical analysis was performed using Kruskal-Wallis test followed by Dunn’s multiple comparisons test, with *P*-values less than 0.05 considered significant. Means are in red. *P*-values are shown only for samples that significantly differ from the control (ds*LacZ*). Data presented are from 4 independent experiments.

## 4. Discussion

To understand the hierarchical organization of CLIP protease cascades and the genetic interactions that control proteolysis within these cascades, we have been using the activation cleavage profiles of cSPHs and cSPs *in vivo* as readouts in genetic epistasis studies. While the cleavage of cSPHs is readily detected in the hemolymph [7, 10, 24], cSP cleavage appears more difficult to probe. This was the case for CLIPB4, CLIPB8 and CLIPB9 in this study, for which no cleavage products were detected, possibly because their active forms are either stably associated with microbial surfaces, since melanization is a highly localized reaction, or rapidly cleared from the hemolymph after inactivation by serpins, which is not the case for cSPHs that lack enzymatic activity and avoid clearance by serpins. The depletion of CLIPB8 from the hemolymph in response to infection suggests that it could be a node in the network on which several proteases converge leading to complete activation.

The mosquito protease cascade bifurcates downstream of CLIPB4 and CLIPB17 into two branches; one converging on CLIPB8 and the other on CLIPB10. A similar observation has been reported in *M. sexta*, whereby two main branches emerge downstream of the cSP HP21, one controls PAP3 activation while the second controls PAP1 in addition to Toll pathway activation [17]. The functional significance of the bifurcation downstream of CLIPB4/B17 remains unclear, especially that CLIPB10 but not CLIPB8 has been shown to function as a PAP [28, 31]. It has been proposed that CLIPB8 functions upstream of CLIPB9, which functions as a PAP, however, CLIPB8 cannot cleave CLIPB9 directly [31].

One of the intriguing observations that emerged from studying these protease cascades in *An. gambiae* is that none of the cSPs involved in melanization has been shown to individually control the activation cleavage of members of the core cSPH module in the hemolymph [7, 24]. Yet, insect cSPs, specifically PAPs, are known to cleave key cSPHs during the melanization response [15, 29, 30]. The fact that CLIPB4, CLIPB9, and CLIPB10 were able to efficiently cleave CLIPA8 *in vitro*, yet when silenced individually they previously failed to rescue the *in vivo* cleavage of CLIPA28, which is just downstream of CLIPA8 in the cSPH module [7], suggests that functional redundancy could mask certain interactions in the context of *in vivo* genetic studies. Based on our data we propose the following updated model of the protease network regulating *An. gambiae* melanization (Fig. S6). Upon infection, TEP1, the most upstream factor in the network [7, 9, 24], triggers the activation of the cSPH module through an unknown mechanism. A yet to be identified modular serine protease (ModSp), in turn, activates the cSP module, the most upstream member of which seems to be CLIPC9 as it controls the activation of CLIPB8 and CLIPB10 (this study), and most likely of CLIPB4 and CLIPB17, although this remains to be verified. The upstream position of CLIPCs with respect to CLIPBs in these cascades has been previously observed in several model insects [3, 4]. CLIPB4 seems to be a central player in the cSP module that directly cleaves multiple substrates, including CLIPA8 of the core cSPH module, CLIPB8 and PPO (this study and [32]). The recent observation that CLIPB4 directly cleaves and activates CLIPB8 *in vitro* [32], further supports our results that CLIPB4 acts upstream of CLIPB8, and strongly suggest that the observed CLIPB8 depletion in the hemolymph after infection is due to its proteolytic processing. We propose that positive reinforcements between cSPs and cSPHs are essential for efficient PPO activation. Initially, systemic infection triggers low-level activation of cSPs, multiple members of which eventually cleave and activate members of the core cSPH module [SPCLIP1-CLIPA8-CLIPA28], as inferred from CLIPA8 cleavage *in vitro*. Active cSPHs, in turn, induce signal amplification, possibly by serving as cofactors or adaptors that may facilitate cSP-cSP and cSP-PPO interactions. The RNAi phenotypes of these core cSPHs characterized by complete abolishment of PO activity and melanin formation [7, 9, 10, 23, 24] support their central roles in the protease network. This is further corroborated by the MelASA phenotype of ds*SPCLIP1* mosquitoes. The role of cSPHs as cofactors for PPO activation is known in other model insects [13, 15, 50], however, our work provides evidence, for the first time, that they also regulate cSP activation by showing that silencing any of the core cSPHs completely rescued CLIPB10 cleavage and CLIPB8 depletion from the hemolymph. The mechanism by which these cSPHs regulate cSP activation and the identity of their direct cSP targets awaits future studies.

Among the cSPs, CLIPB4 and CLIPB17 seem to be central players as they significantly contribute to melanin formation in response to *E. coli* and *M. luteus* infections, however, the contributions of CLIPB8, B9, and B10 were only evident in the context of *M. luteus*, probably because it triggered a more potent melanization response (inferred from the total spot area in Fig. 5), allowing to better gauge the relative contributions of cSPs. In support of this argument, *M. luteus* has been shown to trigger a more potent melanization response than *E. coli* in *Aedes aegypti* [42, 51]. *S. aureus* does not seem to be a good model for MelASA possibly because this bacterium is rapidly and efficiently phagocytosed by hemocytes in mosquitoes [51, 52]. Despite their contributions to melanization, none of the candidate CLIPBs was essential for bacterial clearance from the hemocoel, a process regulated by other effector responses including phagocytosis, antimicrobial peptides and those mediated by TEP1 and the CTL4-CTLMA2 complex [11, 33, 48, 52]. On the other hand, CLIPB14 was the only candidate required for mosquito survival to low dose infections with *E. coli* and *S. aureus*, in agreement with its previously reported role [27]. This is an interesting phenotype, especially because CLIPB14 is not involved in melanization, suggesting that it could be part of the cascade that converges on the activation of the Toll pathway, which remains poorly characterized in mosquitoes.

In conclusion, our work has provided new insight into the structural organization and interaction of cSPs and cSPHs in the protease network that regulates mosquito melanization. The hierarchical structure we propose should guide future studies that employ targeted *in vitro* reconstitution assays and *in vivo* genetic studies to further refine the interactions in the network and unravel the logic behind its complex molecular organization.

## Author Contributions

Conceptualization and study design: Mike A. Osta and Kristin Michel. Experimental work: Sally A. Saab, Xiufeng Zhang, Suheir Zeineddine, Bianca Morejon Viteri. Writing-original draft: Mike A. Osta, Sally A. Saab, and Xiufeng Zhang. Writing-review and editing: Mike A. Osta, Kristin Michel, Xiufeng Zhang and Bianca Morejon.

## Data availability statement

All data generated or analyzed during this study are included in this article. Further inquiries can be directed to the corresponding author.

## Declaration of Competing interest

The authors have no conflicts of interest to declare

## Acknowledgement

We thank Kamal A. Shair Central Research Laboratory for providing free access to their equipment. K.M., X.Z., and B.M. would like to thank all members of the Michel laboratory for mosquito rearing and Dr. Michael Kanost (Kansas State University) for continued support and access to equipment for recombinant protein expression. We also thank Dr. Michael Povelones (University of Pennsylvania) for providing the CLIPC9 antibody.

This work was supported by the National Institutes of Health, National Institute for Allergy and Infectious Disease, grant R01 AI140760 (to K.M. and M.A.O) and by the AUB Research Board award numbers 104107 and 104261 (to M.A.O.). The funding sources had no role in the preparation of data or the manuscript.

## Notes

### Competing Interest Statement

The authors have declared no competing interest.

### Summary of Updates

New supplementary figures S4 and S5 have been added. The abstract has been edited and the manuscript text has been revised to clarify certain statements.

