## Supplemental file for "Insight into the structural hierarchy of the protease cascade that regulates the mosquito melanization response"

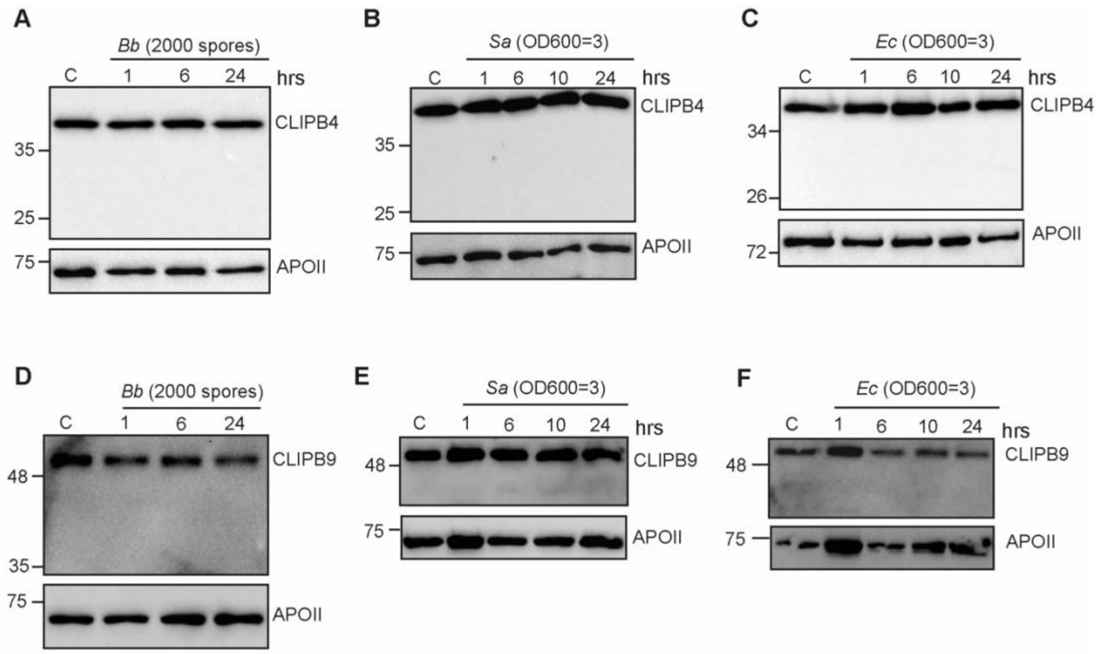


**Fig. S1**. A cleavage product was not detected for CLIPB4 and CLIPB9 after septic infections. (A-C) Western blots of hemolymph extracts obtained at the indicated time points from (A) *B. bassiana* (2000 spores/mosquito), (B) *S. aureus* and (C) *E. coli* injected mosquitoes and probed with αCLIPB4. (D-F) Western blots of hemolymph extracts obtained at the indicated time points from (D) *B. bassiana* (2000 spores/mosquito), (E) *S. aureus* and (C) *E. coli* injected mosquitoes and probed with αCLIPB9. All hemolymph samples were extracted from 40 mosquitoes, quantified using Bradford protein quantification assay and equal amounts of protein were loaded into each well. Membranes were reprobed with αAPOII (without stripping) to control for loading. The figures shown are representative of at least 2 independent biological experiments. *C*, control naïve mosquitoes.

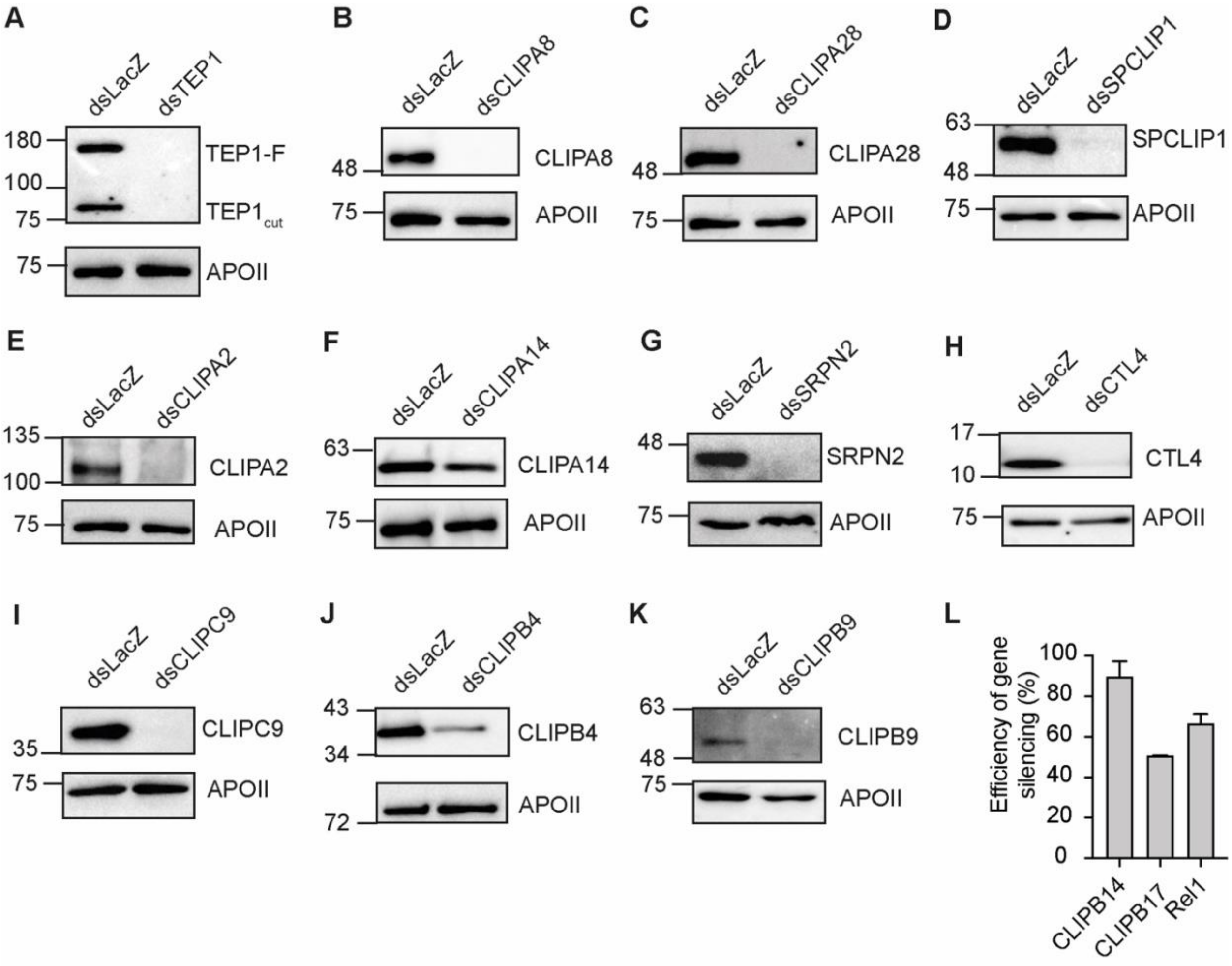


**Fig. S2.** Gene knockdown efficiency by RNAi. (A-K) Western blots showing the gene knockdown efficiency of (A) TEP1, (B) CLIPA8, (C) CLIPA28, (D) SPCLIP1, (E) CLIPA2, (F) CLIPA14, (G) SRPN2, (H) CTL4, (I) CLIPC9, (J) CLIPB4, and (K) CLIPB9 in naïve mosquitoes at 3 days post dsRNA injection. All hemolymph samples were extracted from 40 mosquitoes, quantified using Bradford protein quantification assay and equal amounts of protein were loaded into each well. Membranes were reprobed with αAPOII as loading control. (L) Efficiency of gene silencing of CLIPB14, CLIPB17 and Rel1 in naïve mosquitoes at 3 days after dsRNA injection as measured by qRT-PCR. Data shown here are from 2 independent trials. Error bars represent standard error of the mean.

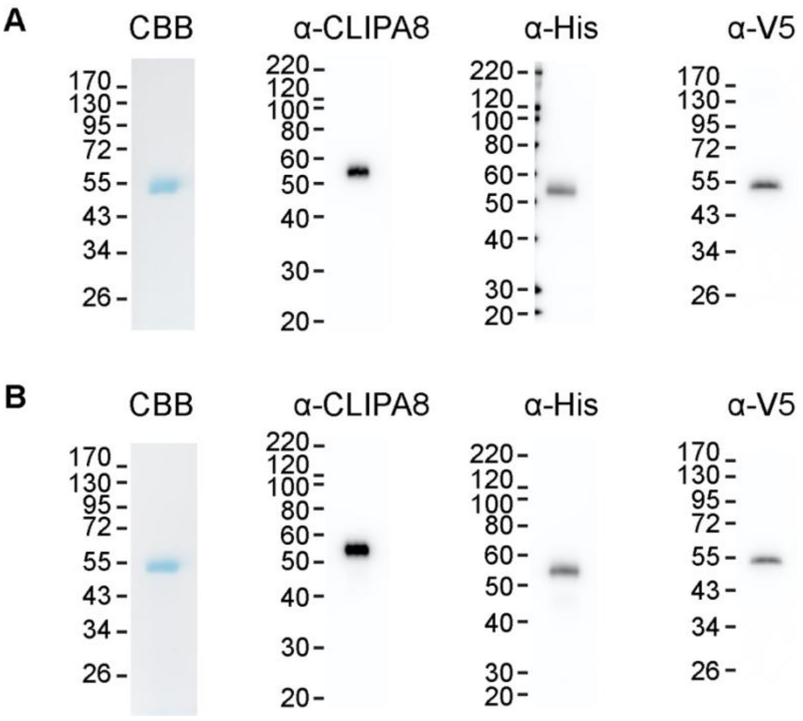


**Fig. S3.** Purified recombinant CLIPA8 and CLIPA8$_{Xa}$. After the purification of recombinant proCLIPA8-V5-His (A) and proCLIPA8$_{Xa}$-V5-His (B), proteins were subjected to SDS Page and detected by Coomassie Brilliant blue (CBB) staining (1 µg, amount of examined protein), or further submitted to western blot and detected by anti-CLIPA8 antibody (10 ng), anti-His antibody (30 ng) and anti-V5 antibody (10 ng). Recombinant proCLIPA8 and proCLIPA8$_{Xa}$ were detected at ~54 KDa with a high purity.

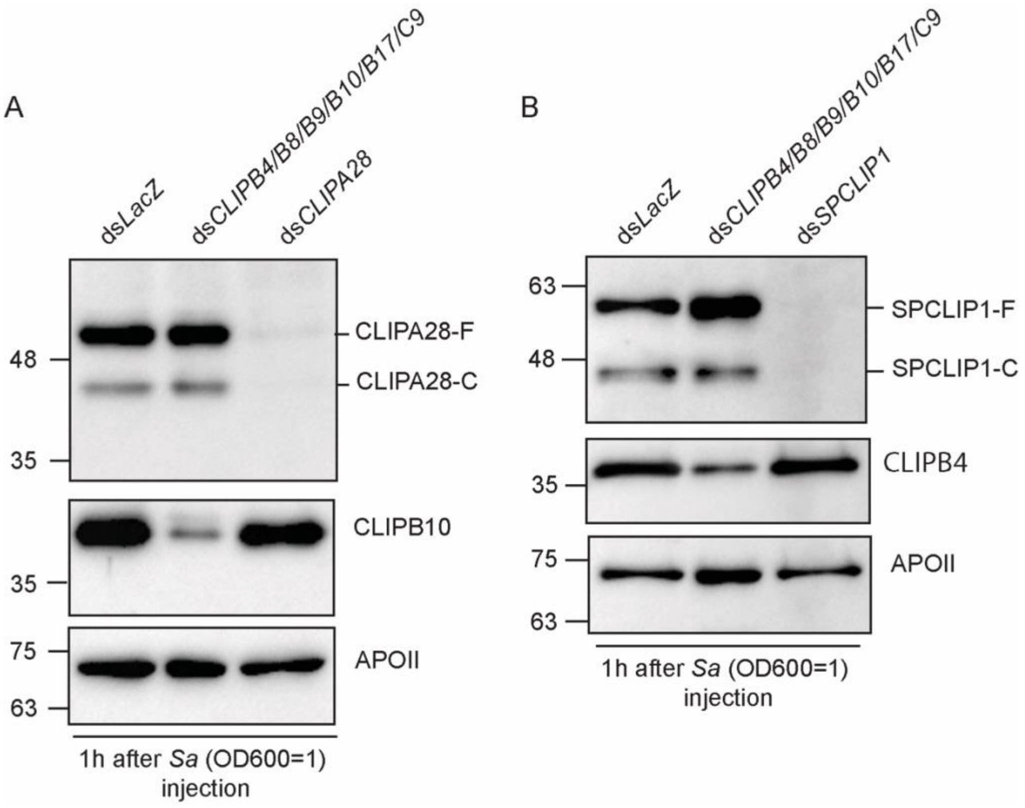


**Fig. S4.** The simultaneous knockdown of 6 key cSPs does not abolish the cleavage of CLIPA28 nor SPCLIP1. (A-B) Western blots of hemolymph extracts obtained at 1 hour after *S. aureus* (*Sa*) (OD600=1) injection of the indicated mosquito genotypes and probed with (A) αCLIPA28 and (B) αSPCLIP1. Membranes were reprobed with (A) αCLIPB10 and (B) αCLIPB4 to show that the simultaneous introduction of multiple gene-specific dsRNA species does not influence the efficiency of gene knockdown. Membranes were also reprobed with αAPOII as loading control.

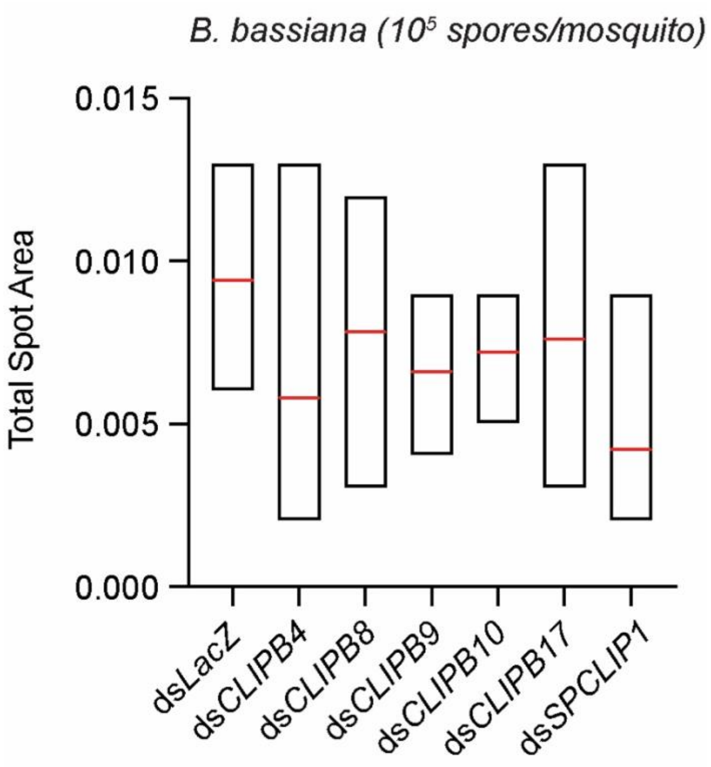


**Fig. S5.** The candidate CLIPBs and SPCLIP1 do not significantly contribute to *B. bassiana* spore melanization. MelASA assay conducted on the indicated mosquito genotypes at 14 hrs post *B. bassiana* spore injection (100000 spores/mosquito). The amount of melanin deposits present in infected mosquito excreta was measured from Whatman papers placed at the bottom of the paper cups housing 35 mosquitoes per sample. Statistical analysis was done using One-Way ANOVA with Dunnett's post-test to compare multiple treatment groups. Means are shown in red lines. Data shown are from 5 independent trials.

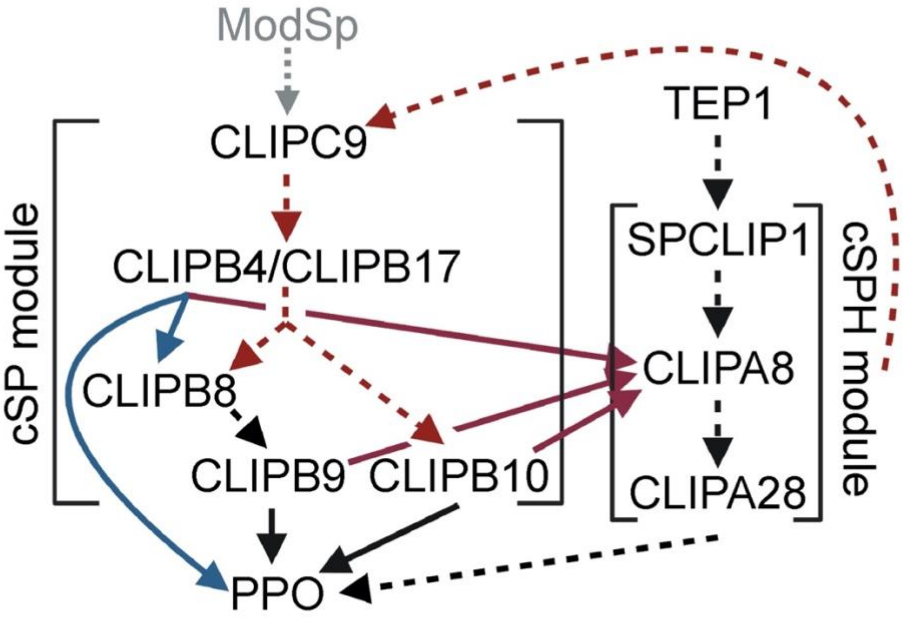


Fig. S6. Schematic drawing of the protease network regulating mosquito melanization. The scheme indicates the current knowledge of hierarchy in the cSP and cSPH modules constituting the network. Dashed lines indicate hierarchy inferred from genetic studies *in vivo* (i.e. hemolymph) described in this study (red) or previous work (black; [1-4]). Solid lines indicate direct cleavage events inferred from *in vitro* assays described in this study (red), previous work (black; [5, 6]) or in the recent study of Zhang *et al*. (blue; [7]). Dashed, blue line represents the presumably promiscuous cleavage of members of the cSPH modules by several cSPs. The module serine protease (ModSp) that acts upstream of cSPs is shown in grey as its identity is still unknown in *An. gambiae*. Dashed lines indicate that the enzymatic steps are not yet fully characterized.

### Table S1: Primers used for dsRNA production

| Gene | Primers used for dsRNA synthesis (T7 promoter sequence underlined; 5'-3') | Reference |
|---|---|---|
| *LacZ* | For: TAATACGACTCACTATAGGGAGAATCCGACGGGTTGTTACT<br>Rev: TAATACGACTCACTATAGGGCACCACGCTCATCGATAATTT | [8] |
| *TEP1* (AGAP010815) | For: TAATACGACTCACTATAGGGTTTGTGGGCCTTAAAGCGCTG<br>Rev: TAATACGACTCACTATAGGGACCACGTAACCGCTCGGTAAG | [9] |
| *SPCLIP1* (AGAP028725) | For: TAATACGACTCACTATAGGGGTCACCGAACACGGCCAAC<br>Rev: TAATACGACTCACTATAGGGATCGAAGCTGATCGGATCGGG | [2] |
| *CLIPA8* (AGAP010731) | For: TAATACGACTCACTATAGGGAACAACGAACCCGTAGAATATG<br>Rev: TAATACGACTCACTATAGGGGGTTAGCGCCTCGATACC | [10] |
| *CLIPA28* (AGAP010730) | For: TAATACGACTCACTATAGGGAGACCACCAAGGAACCGTTCCCGCA GCAA<br>Rev: TAATACGACTCACTATAGGGAGACCGCAACCGATGCCCCACGAT ACGAT | [1] |
| *SRPN2* (AGAP006911) | For: TAATACGACTCACTATAGGGCTGGTCAATGTGATCTACTT<br>Rev: TAATACGACTCACTATAGGGATTGTTCCGAGGGTTTCAT | [1] |
| *CLIPC9* (AGAP004719) | For: TAATACGACTCACTATAGGGGGTGCAGTAAGAAGGCCCAT<br>Rev: TAATACGACTCACTATAGGGACTGCATGTCCAAGCAATCC | [3] |
| *CLIPA2* (AGAP011790) | For: TAATACGACTCACTATAGGGATCCTAACAACGGCACACTGTGTGA<br>Rev: TAATACGACTCACTATAGGGTCCTGATCGCCATGATTGGTGGTGCT | [11] |
| *CLIPA14* (AGAP011788) | For: 5'- TAATACGACTCACTATAGGGCGGCATCATCGACATCCGTGTC-3'<br>Rev: 5'- TAATACGACTCACTATAGGGGTTGCTGTCGGCGACACGCTCCT-3' | [12] |
| *CLIPB4* (AGAP003250) | For: TAATACGACTCACTATAGGGAGTAGCGGTCGTGCATCAGA<br>Rev: TAATACGACTCACTATAGGGTGGCCTGCTAGAGCCAGCGT | [1] |
| *CLIPB8* (AGAP003057) | For: TAATACGACTCACTATAGGGGTCATACCGCACCCGGAGTA<br>Rev: TAATACGACTCACTATAGGGTTCCCGTTCGACGTACGGCA | [1] |
| *CLIPB9* (AGAP029769) | For: TAATACGACTCACTATAGGGAATGCACGACACCGACGAGGT<br>Rev: TAATACGACTCACTATAGGGGTTTGCCCTCCTTGCGCTCA | [1] |
| *CLIPB10* (AGAP029770) | For: TAATACGACTCACTATAGGGGAGCGTAAGGGATGAGTTCT<br>Rev: TAATACGACTCACTATAGGGCAGCACGTACCGTCCGTTGA | [1] |
| *CLIPB14* (AGAP010833) | For: TAATACGACTCACTATAGGGGACTGCAAGCAGGTCAAAGGC<br>Rev: TAATACGACTCACTATAGGGTCCACGGAACATCTCCCGCT | [1] |
| *CLIPB17* (AGAP001648) | For: TAATACGACTCACTATAGGGAGCGTGGGGAATTCCCGTGGA<br>Rev: TAATACGACTCACTATAGGGGGATCGTCCATCAGCAGCGA | [1] |
| **Primers used for dsRNA production in the *M. luteus* MelASA that are different from those above** (T7 promoter sequence underlined; 5'-3') | | |
| *GFP* | For: TAATACGACTCACTATAGGGCGATGC<br>Rev: TAATACGACTCACTATAGGGCGGACT | [5] |
| *CLIPB4* (AGAP003250) | For: TAATACGACTCACTATAGGGTCAGGATTGCGTGAATCCGG<br>Rev: TAATACGACTCACTATAGGGAGTTTTACGCTCGTAGAGAC | |
| *CLIPB8* (AGAP003057) | For: TAATACGACTCACTATAGGGTGTGCGACATCCCGAACGAG<br>Rev: TAATACGACTCACTATAGGGCATCCGGTCCGAGATTGTCC | [4] |
| *CLIPB10* (AGAP029770) | For: TAATACGACTCACTATAGGGCCGCAAAACTTTGGCATC<br>Rev: TAATACGACTCACTATAGGGCTGGTAGGCGGGAAGTCAT | [6] |
| *CLIPB14* (AGAP010833) | For: TAATACGACTCACTATAGGGATGTCGTACCAGCTGAAGCA<br>Rev: TAATACGACTCACTATAGGGCTACGCCAAAATTCCACGGA | |

**Table S2: Primers used for qRT-PCR**

| Gene | (5'-3') | Reference |
|---|---|---|
| *AgS7* | For: AGAACCAGCAGACCACCATC<br>Rev: GCTGCAA ACTTCGGCTATTC | [13] |
| *Rel1*<br>(AGAP009515) | For: CCAACCTCGATCCGGTGTTCA<br>Rev: TAGGTCGGTCGTGGAAAGTGA | [14] |
| *CLIPB14*<br>(AGAP010833) | For: GTTAGCAAGCGGTTTGTGCT<br>Rev: TGAAATTCCATTCCGCAACGC | [1] |
| *CLIPB17*<br>(AGAP001648) | For: TGCACGACCGAAGGCATTA<br>Rev: AGCTTTTTGTGCGTGGCAC | [1] |

**Table S3: Primers used for CLIPA8 protein expression constructs**

| | |
|---|---|
| **Synthesized BglII-V5-6His-PacI fragment** | |
| 5'- GGAGATCTAGGTAAGCCTATCCCTAACCCTCTCCTCGGTCTCGATTCTACGCGTACCGGT*CATCATCACCATCACCAT*TGATTAATTAATA -3'<br>5' BglII and 3' PacI cloning sites are underlined. 6His tag sequence is italized. V5 tag sequence is double-underlined. | |
| **Amplification of full-length CLIPA8 with restriction sites into pOET3-V5-His vector** | |
| CLIPA8-F-NotI | 5'-ATGCGGCCGCATGCCTAGCTGGTGGTGT-3' |
| CLIPA8-R-BglII | 5'-CCTAGATCTCCTCCTCCCAGTATTCCCTCGATTGT-3' |
| Restriction sites are underlined. | |
| **CLIPA8 mutagenesis** | |
| CLIPA8mutag | 5'- CGAGGCGAATGCGCAGAACCAGGAGATCATTGAAGGACGCTTCGGCGAGGAAG-3' |
| Sequence encoding IEGR is underlined. | |
